# Inference of ploidy by leveraging read depth from amplicon sequencing

**DOI:** 10.1101/2020.07.30.229500

**Authors:** Thomas A. Delomas, Stuart C. Willis, Andrea Schreier, Shawn Narum

## Abstract

Variation in ploidy occurs naturally in select plant and animal species. Ploidy variation can also occur spontaneously or be induced during artificial propagation of fish and shellfish. Studying species and systems that have variable ploidy requires techniques to infer ploidy of individuals. Massively parallel sequencing of biallelic SNPs has been used to infer ploidy, but existing techniques have several drawbacks. These include being limited to only comparing a fixed number of ploidies (diploidy, triploidy, and tetraploidy) and requiring that heterozygous genotypes in an individual be identified prior to ploidy inference. We describe a method of inferring ploidy from sequencing of biallelic SNPs based on beta-binomial mixture models. This method is generalized to apply to any ploidy and does not require prior identification of heterozygous genotypes. We demonstrate efficacy of this method for comparing ancestral octoploidy, decaploidy, and dodecaploidy (tetraploidy, pentaploidy, and hexaploidy for the sequenced SNPs) in white sturgeon and diploidy and triploidy in Chinook salmon with amplicon sequencing (GT-seq) data. Results indicated that ploidy could be reliably estimated for individuals based on distinct distribution of log-likelihood ratios (LLR) for known ploidy samples of both species that were tested. Confidence in ploidy estimates increased with sequencing depth. We encourage users to explore the sequencing depths and LLR critical values that provide reliable estimates of ploidy for a given organism and set of SNPs. We expect that the R package provided will empower studies of genetic variation and inheritance in organisms that vary in ploidy naturally or as a result of artificial propagation practices.

## Introduction

The number of haploid copies of the genome in somatic cells of an individual, termed ploidy, is naturally variable in some plant and animal species (Lamatsch & Stöck, 2009; Mock et al., 2012; Yamashita, Jiang, Onozato, Nakanishi, & Nagahama, 1993; Zhang & Arai, 1999). Ploidy variation in fish has been observed to occur spontaneously (Gold & Avise, 1976; Machado et al., 2012; Utsunomia et al., 2014). Some artificial spawning and rearing practices may increase the rate of spontaneous ploidy variation (Aegerter & Jalabert, 2004; Cherfas, Gomelsky, Ben-Dom, & Hulata, 1995; Delomas & Dabrowski, 2016; Flajšhans, Kvasnička, & Ráb, 1993; Glover et al., 2015; Thorgaard et al., 1982; Van Eenennaam et al., 2020) which could lead to more ploidy variation in systems stocked with hatchery-origin fish. Alterations in ploidy can also be induced in fish and shellfish to yield individuals with advantageous qualities for cultivation, such as sterility (Benfey, 1999; Nell, 2002), or stocking in water bodies where reproduction of stocked fish is undesirable (Cassinelli, Meyer, Koenig, Vu, & Campbell, 2018). Ploidy variation is linked to reproductive isolation (Husband & Sabara, 2004; Husband, Schemske, Burton, & Goodwillie, 2002), speciation (Ptacek, Gerhardt, & Sage, 1994; Wood et al., 2009), changes in gamete ploidy and reduced fertility (Delomas & Dabrowski, 2018; Feindel, Benfey, & Trippel, 2010; Liu et al., 2001), and differences in metabolism (Hyndman, Kieffer, & Benfey, 2003; Leal, Clark, Van Eenennaam, Schreier, & Todgham, 2018). Studies in systems and species with variable ploidy that do not account for such variation therefore risk ignoring an important confounding factor.

Techniques to infer the ploidy of individual samples are required when the study system and species have variable ploidy. Ploidy is commonly inferred using flow cytometry to directly measure nuclear DNA content in cells from a blood or solid tissue sample (Delomas & Dabrowski, 2018). A Coulter counter is also commonly used in animals when a fresh blood sample can be easily obtained (Wattendorf, 1986). However, these techniques can be untenable for ploidy determination when sampling is conducted in remote locations, far from a flow cytometer, Coulter counter, or the reagents and consumables needed to fix samples for future flow cytometry. A researcher may also wish to determine the ploidy of an archived tissue sample, such as from a museum specimen. Because fresh or specially fixed tissues are required for Coulter counter or flow cytometry analyses, another method is necessary to determine ploidy of archived samples.

To address this shortcoming, methods have been developed to infer ploidy from massively parallel sequencing data, namely read counts at biallelic single nucleotide polymorphisms (SNPs). One graphical technique assisted by ploidyNGS (Augusto Corrêa dos Santos, Goldman, & Riaño-Pachón, 2017) is to visually inspect histograms of allele depth ratios (Figure 1). The number and location of peaks corresponding to heterozygous genotypes can be used to classify samples. A drawback of this technique is that it requires visual, not statistical, evaluation of histograms.

**Figure 1.**
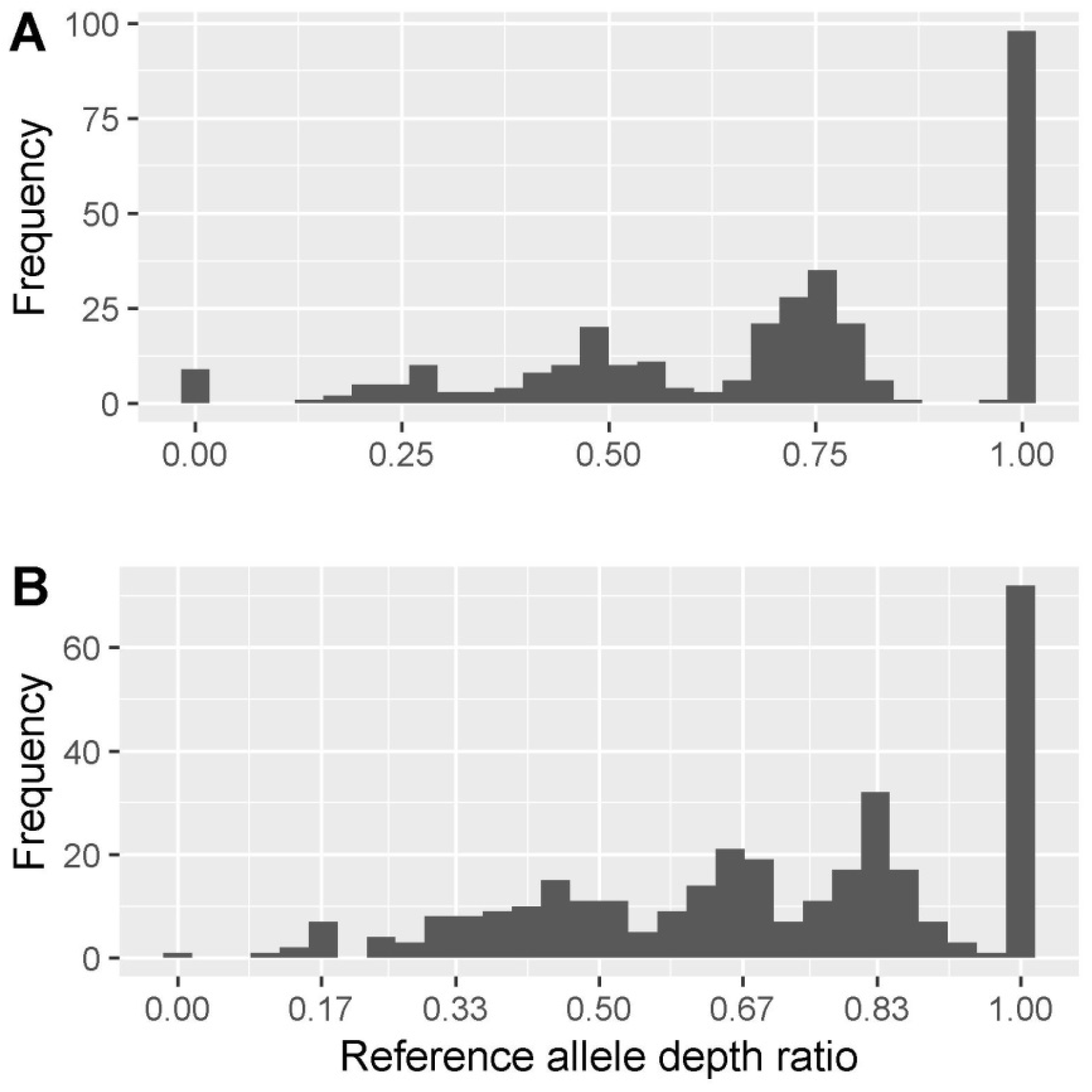
Distribution of the reference allele depth ratio (depth of reference allele / total depth) for two white sturgeon of known ploidy. These graphs were generated similarly to the method used by ploidyNGS (Augusto Corrêa dos Santos et al., 2017) A) True ancestral octoploid (tetraploid for the genotyped SNPs) B) True ancestral dodecaploid (hexaploid for the genotyped SNPs)

Several methods that allow more automated ploidy inference have been developed. The R package gbs2ploidy (Gompert & Mock, 2017) uses read counts to estimate relative allele dosage at heterozygous loci. These estimates and observed heterozygosity are then used to categorize sample ploidy with clustering algorithms. One drawback of this method is that it requires prior identification of heterozygous genotypes. At higher levels of ploidy (e.g. octoploidy), confidently separating homozygous genotypes from genotypes with only one copy of the minor allele can require a large number of reads. In some situations, the depth required may be difficult to achieve across a sufficient number of loci due to low sample quality or cost constraints. Additionally, the current implementation of gbs2ploidy is limited to only discriminating between diploidy, triploidy, and tetraploidy.

The program nQuire (Weiß, Pais, Cano, Kamoun, & Burbano, 2018) models observed ratios of allele depth at heterozygous SNPs with a Gaussian mixture model. The means of the Gaussians correspond to the allele dosage expected with a given ploidy (e.g., 1/3 and 2/3 for triploidy). The use of Gaussian distributions, as compared to binomials, allows higher levels of dispersion in the data to be modeled. The authors additionally demonstrate that a uniform noise component can be added to model spurious observations. However, a drawback of this approach is that modelling the ratios of allele depth, and not the read counts, ignores the relationship between variance and depth. Second, the method requires identification of loci with heterozygous genotypes in each individual as homozygous genotypes are not modelled. As mentioned above, confidently separating homozygous genotypes from genotypes with one copy of the minor allele can require a prohibitive number of reads at higher levels of ploidy. Third, this approach is currently only implemented in nQuire for discriminating between diploidy, triploidy, and tetraploidy.

A method based on a likelihood ratio statistic was developed to address some of the shortcomings of the above approaches, as well as account for variable sequencing error and allelic bias between loci (Delomas, 2019). This method excludes homozygous loci using a binomial test and then calculates a likelihood ratio comparing diploidy and triploidy. The likelihoods are calculated by assuming the read counts are binomial random variables. One drawback of this method is that it is limited to comparing diploidy and triploidy. Additionally, while it does not require the user to identify heterozygous loci, it attempts to identify them using a binomial test. A final drawback is that modelling the read counts as binomials does not allow for overdispersion which can be present in some sequencing data. As demonstrated (Delomas, 2019), this method performs well for differentiating diploids and triploids with amplicon sequencing data, but a similar strategy may not be suitable for differentiating ploidy levels with more similar allele dosages.

Our goal was to develop a method of inferring ploidy from high throughput sequencing data for biallelic SNPs that addressed the drawbacks of existing methods and was applicable to any ploidy. Our motivation stems from the case of white sturgeon (*Acipenser transmontanus*). White sturgeon are ancestral octoploids (8n) (Drauch Schreier, Gille, Mahardja, & May, 2011), but spontaneous autopolyploidy has been observed in hatchery settings, producing dodecaploids (12n) (Van Eenennaam et al., 2020). Crossing individuals with these two ploidy levels then yields decaploids (10n). Although flow cytometry and Coulter counter analysis can be used to accurately distinguish between white sturgeon of different ploidies (Fiske et al., 2019), these techniques cannot be used for archived tissue samples. However, a panel of biallelic SNPs was developed (Willis et al., 2020) that are detected in four copies in the genomes of the ancestrally octoploid white sturgeon. We developed a method to efficiently distinguish between tetraploidy, pentaploidy, and hexaploidy using these SNPs, corresponding to octoploidy, decaploidy, and dodecaploidy on the ancestral scale, respectively. We here describe the method and validate it both with white sturgeon of three ploidies and Chinook salmon of two ploidies.

## Methods

### Beta-binomial mixture model

For the purpose of this explanation, we use the term “reference allele” to refer arbitrarily to one of the alleles of a biallelic SNP. A biallelic SNP in an individual with ploidy *x* has *x* + 1 possible states corresponding to 0, 1,…, *x* copies of the reference allele. As such, we model counts of the reference allele as a mixture with *x* + 1 components. The weights of the components correspond to the proportion of SNPs in each state. A similar approach is implemented in nQuire except that components corresponding to states 0 and *x* are not included (Weiß et al., 2018).

Observation of a read of the reference allele can be considered a Bernoulli random variable with probability of success dependent upon the true state of the SNP, *x*, the rate of sequencing error, and allelic bias. Gerard et al. (2018) derived an equation for calculating this probability. Because individual reads can be considered Bernoulli random variables, it is natural to model the read count as a binomial random variable. Considering a mixture of binomials, the likelihood of observing *c_i_* counts of the reference allele given *n_i_* total reads and a ploidy of *x* is

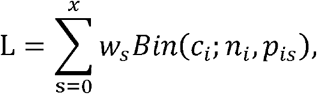

where *w_s_* is the weight of the component for state *s* and *p_is_* is the probability of observing a read of the reference allele for locus *i* and state *s* calculated according to Gerard et al. (2018).

Overdispersion is commonly present in sequencing data. One solution is to model read counts with a beta-binomial distribution and an overdispersion parameter, τ (Gerard et al., 2018). We adopt this approach, and the likelihood now becomes

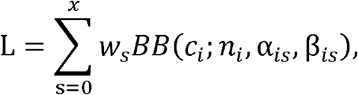

with

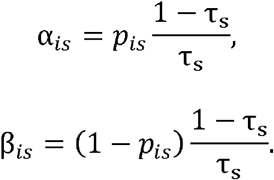

Gerard et al. (2018) developed a method to call genotypes in groups of individuals of known ploidy and inferred values of τ that were specific to each locus. In the current application, we wanted to avoid using information from multiple individuals to simplify application of this method and to eliminate the need to either assume that τ is constant across ploidies or develop a mechanism for estimating different values of τ for each possible ploidy using samples of unknown ploidy. Therefore, a value of τ was defined for each state, τ_*s*_, within a given individual and ploidy. This decision implies that variance depends on the true state of a locus. nQuire similarly fits the variance of Gaussian distributions separately between states within an individual (Weiß et al., 2018). Assuming independence between loci, the total likelihood is the product of the likelihoods of each locus.

Addition of a uniform noise component to the Gaussian mixture model used by nQuire was demonstrated to be helpful when analyzing noisy data (Weiß et al., 2018). The same addition is possible with a beta-binomial mixture model. The model then has *x* + 2 components and the likelihood for one locus is

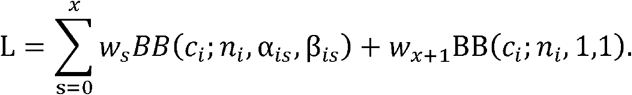

This model can be fit using an expectation-maximization (EM) algorithm. In the expectation step, weights of each component are updated as typical for a mixture model

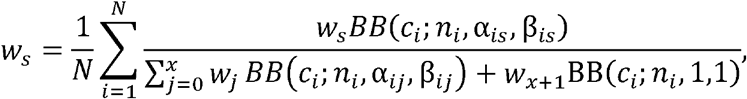

with all elements of *w* being updated using the values of *w* from the previous iteration. The maximization step updates the values of τ by maximizing the log-likelihood using the method of Byrd et al. (1995) informed by the analytical gradient.

Functions to fit this model as well as the intermediary models described (binomial mixture model and beta-binomial without uniform noise) are implemented in an R package, tripsAndDipR v0.2.0, available at www.github.com/delomast/tripsAndDipR.

### Inferring ploidy

To infer ploidy of an individual, models with different ploidies need to be compared. The maximum likelihood estimate (MLE) is the ploidy corresponding to the model with the highest likelihood. Models can be more quantitatively compared by calculating a log-likelihood ratio between two models. If more than two ploidies are possible, a series of log-likelihood ratios (LLR) comparing the most likely model to all others can be calculated. Rejection of less likely models depends on the distribution of the log-likelihood ratios. These distributions are unknown but can be empirically approximated by assessing samples of known ploidy. If a reliable method of simulating data given ploidy is available for a particular sequencing protocol, a Monte Carlo method for approximating these distributions could also be used.

### Assessing the method

To assess the efficacy of this method, we applied it to three groups of samples. All samples were fin clips. The first group, subsequently referred to as “known ploidy sturgeon”, consisted of 19 octoploid and 23 dodecaploid white sturgeon. These sturgeon were from a Central California caviar farm (original broodstock source: Sacramento River). Ploidy was confirmed using a Coulter counter at the University of California Davis. These samples were genotyped at 325 SNPs according to Willis et al. (2020). Samples were initially sequenced targeting a depth ten times higher than Willis et al. (2020) recommended to achieve high genotyping success, and reads were then randomly down-sampled at levels of 10%, 30%, and 50% to evaluate the effect of sequencing depth on ploidy inference.

The second group, subsequently referred to as “presumed decaploid sturgeon”, consisted of 17 full-sibling white sturgeon produced at a caviar farm in Central California (broodstock source: Sacramento River). Individual parents had been identified as octoploid and dodecaploid through the method described here, and so the method was applied to their offspring who were presumed to be decaploid. This group of sturgeon was genotyped according to Willis et al. (2020). In both groups of sturgeon, tetraploid, pentaploid, and hexaploid models were considered.

The third group of samples, subsequently referred to as “known ploidy salmon”, consisted of 93 triploid Chinook salmon *Oncorhynchus tshawytscha* from Idaho Department of Fish and Game’s Nampa Fish Hatchery whose ploidy was confirmed by flow cytometry and 80 diploid Chinook salmon from Idaho Department of Fish and Game’s Rapid River Fish Hatchery (diploidy confirmed by successful reproduction). Fin clips were taken from these fish and genotyped according to the GT-seq method of amplicon sequencing (Campbell, Harmon, & Narum, 2015) with a panel of 342 SNPs. Diploid and triploid models were considered for these samples.

We assumed no allelic bias and sequencing error rates of 0.01 (1%) for all loci in all analyses.

## Results

In the known ploidy sturgeon, mean depth of SNPs within individuals had mean ± SD of 528 ± 249, 1585 ± 746, 2642 ± 1243, 5283 ± 2486 reads per locus at subsampling levels of 10, 30, 50, and 100%, respectively. Mean depth of SNPs within individuals of the presumed decaploid sturgeon and known ploidy salmon was 752 ± 284 and 300 ± 92, respectively. The MLEs for the known ploidy sturgeon in all subsampling levels and the known ploidy salmon were correct. The MLEs for the presumed decaploid sturgeon were all decaploid, fitting expectations based on the inferred ploidy of their parents. The LLRs comparing the true ploidy with alternative ploidies were centered away from zero (Figures 2, 3, and 4) and the distance increased with increasing depth (Figure 2).

**Figure 2.**
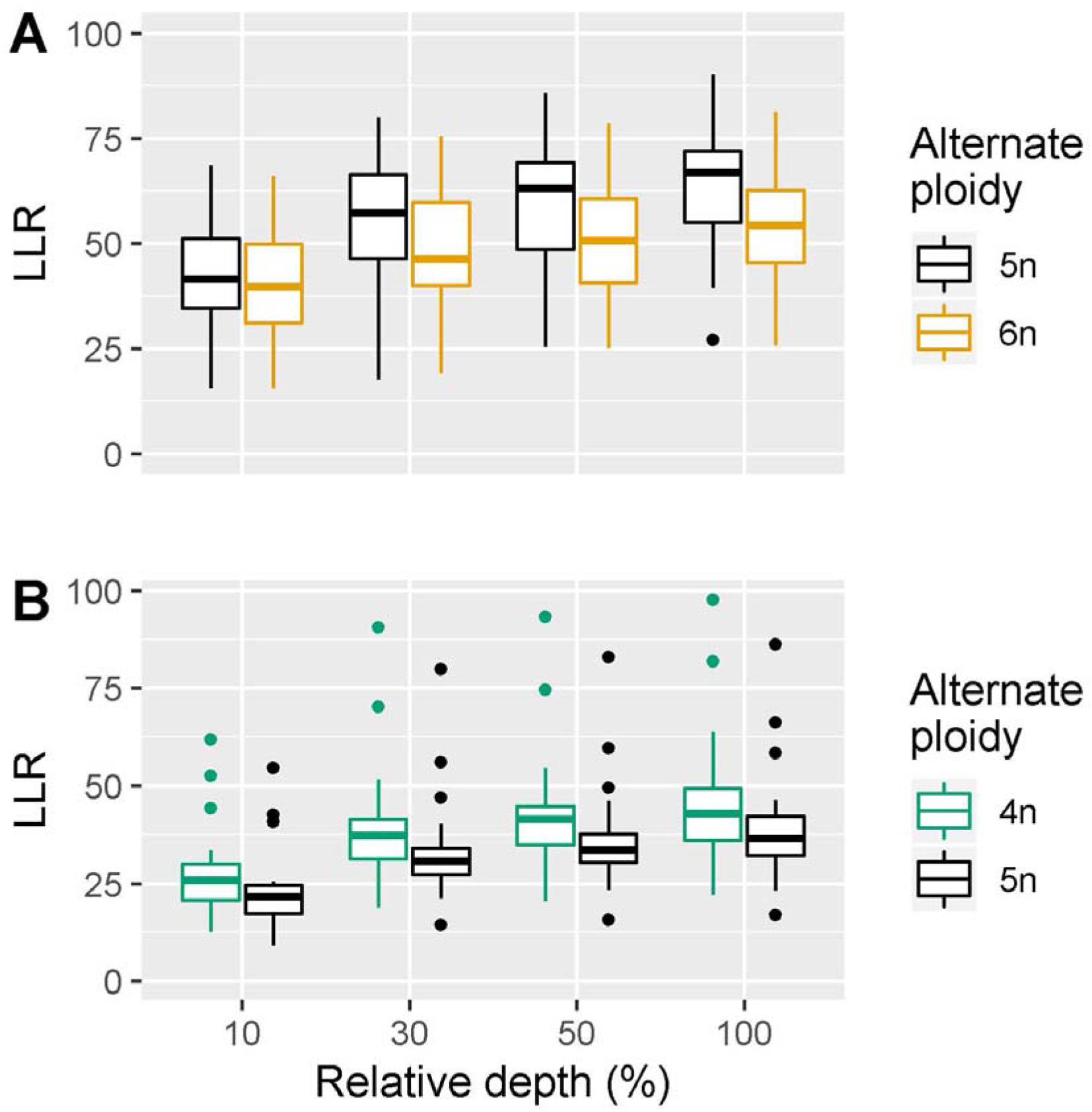
Boxplots of LLR for known ploidy white sturgeon at various depths. Values of LLR are comparing the true ploidy with the alternate ploidy. 4n, 5n, 6n represent tetraploid, pentaploid, and hexaploid, respectively A) True ancestral octoploids (tetraploid for the genotyped SNPs) B) True ancestral dodecaploids (hexaploid for the genotyped SNPs)

**Figure 3.**
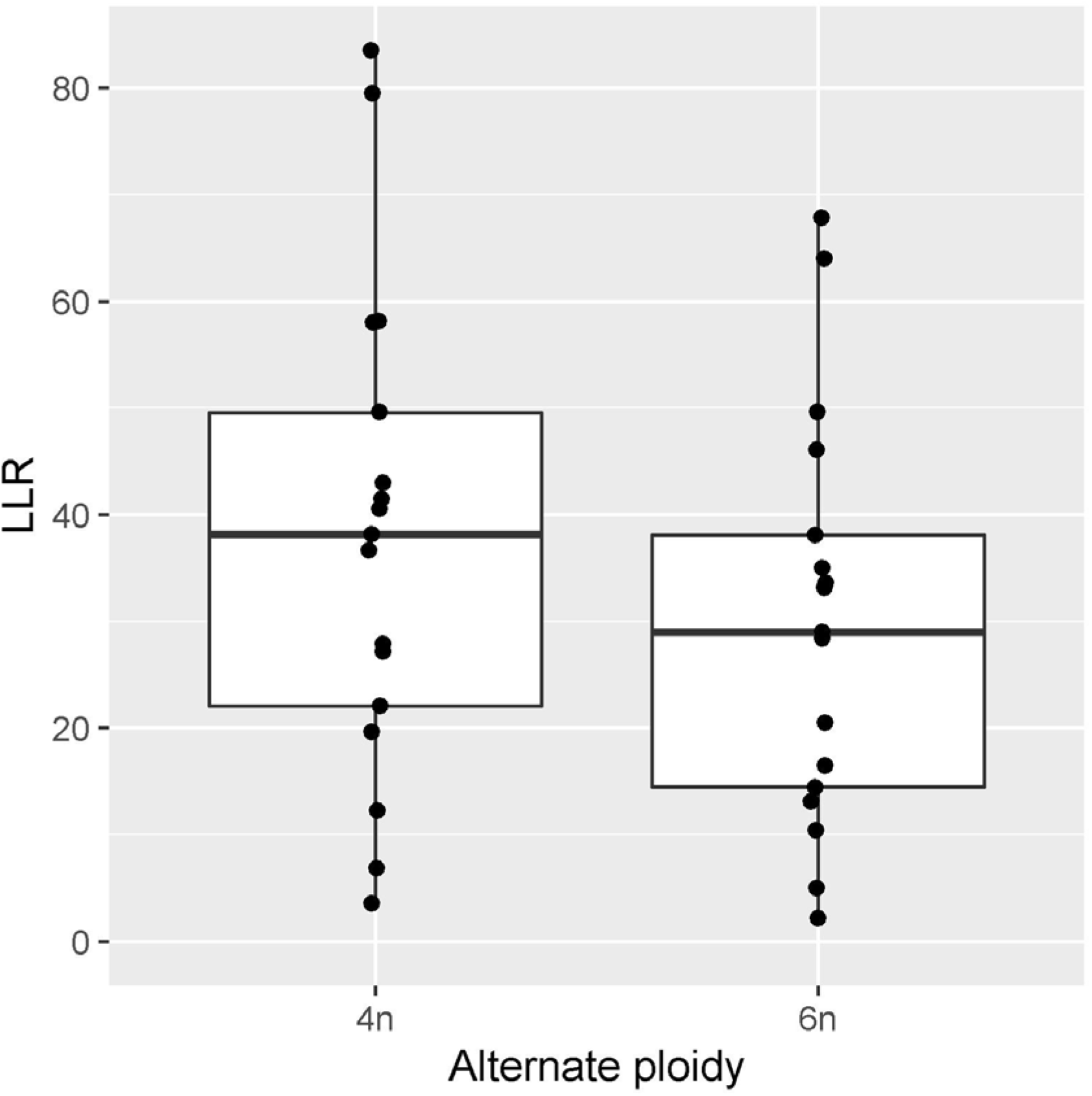
Boxplots of LLR for ancestral decaploid (pentaploid for the genotyped SNPs) white sturgeon. Values of LLR are comparing pentaploidy with the alternate ploidy. All data points are plotted overlying the boxplots. 4n and 6n represent tetraploid and hexaploid, respectively.

**Figure 4.**
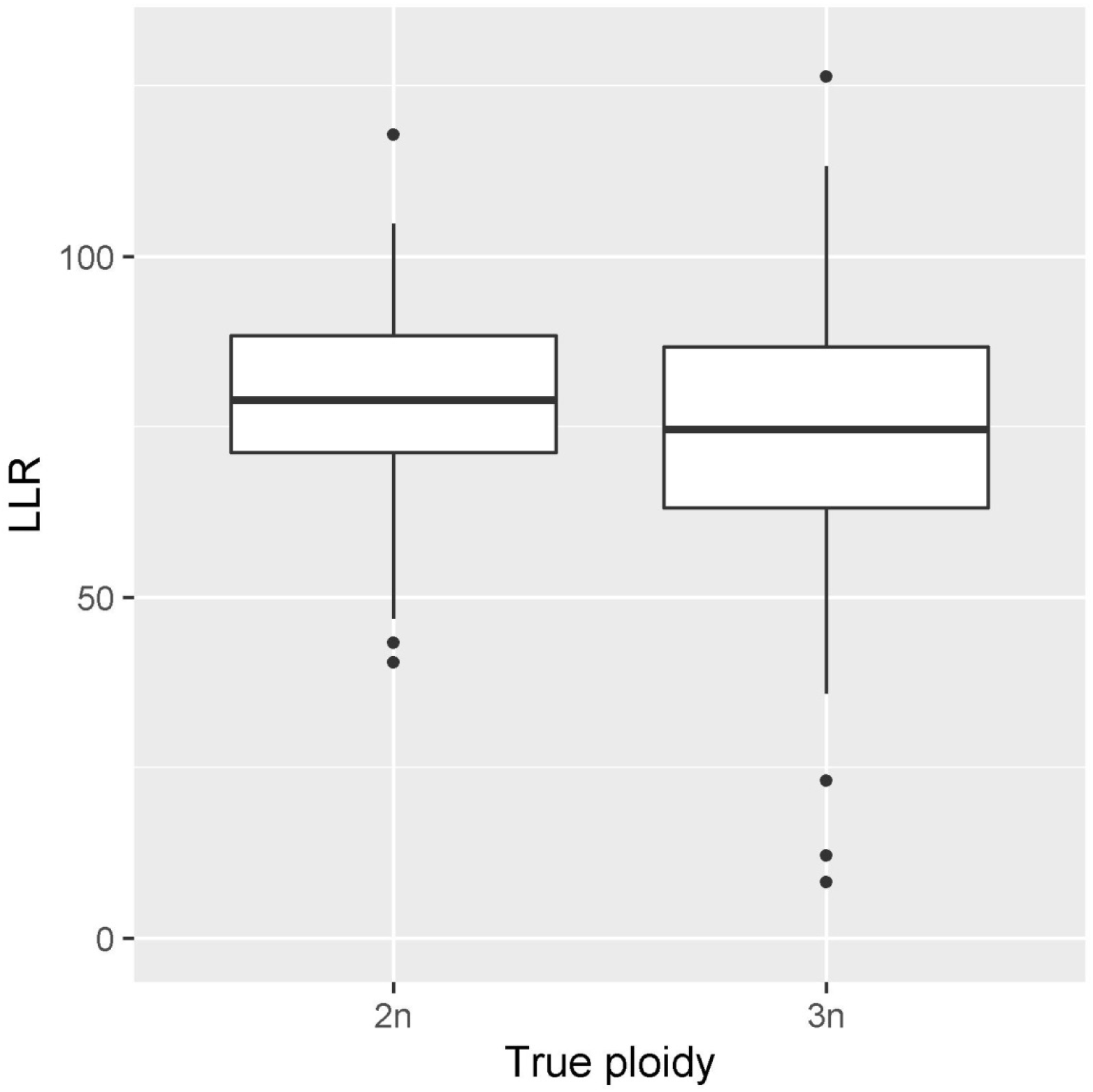
Boxplots of LLR for known ploidy Chinook salmon. Values of LLR are comparing the true ploidy (x-axis) with the opposing ploidy. 2n and 3n represent diploid and triploid, respectively

Fitting the described beta-binomial model and comparing ploidies through LLR accurately separated individuals according to true ploidy. The magnitude of separation of different ploidies with a given set of SNPs was dependent on sequencing depth (Figure 2). The lowest down-sampling level (10%), which corresponds to targeting the depth recommended for genotyping by Willis et al. (2020), gave accurate MLEs for ploidy. While not demonstrated in these analyses, the statistical model implied that the magnitude of separation was also influenced by the variability of the SNPs in the sequenced individuals. Genotypes with relative allele dosages that were shared between ploidies did not contribute information about ploidy. This included homozygous genotypes and, in comparisons of tetraploidy and hexaploidy, genotypes with equal numbers of both alleles (relative dosage of 0.5).

We found that the distribution of LLRs for white sturgeon samples of known and presumed ploidy could be used to set critical values for rejecting less likely models. With the panel of SNPs and depth targeted in this study, a critical value of 10 was appropriate for rejecting alternative ploidy models. Very few individuals had an LLR less than 10, and this critical value did not result in any false classifications of the known ploidy samples (Figures 2 and 3).

## Discussion

We describe here a new statistical model for inferring ploidy from sequencing data and demonstrate its efficacy using amplicon sequencing for two species of varying ploidy levels. Increased sequencing depth increased the likelihood of the correct ploidy (Figure 2). The relationship between mean depth and accuracy of inferred ploidy depends on the ploidies being assessed, the number and variability of loci, and the desired level of confidence. Additionally, the observed variance in the read counts over what would be expected from a binomial random variable (overdispersion) impacts the depth required. As such, we recommend users evaluate minimum mean depth requirements for their panel and species.

Unlike previous methods (Delomas, 2019; Gompert & Mock, 2017; Weiß et al., 2018), the current method is generalized to assess any ploidy and does not require identification of heterozygous genotypes in an individual prior to ploidy inference. However, as noted by Gompert and Mock (2017), inferring ploidy from sequencing data cannot separate individuals of lower and higher ploidy when the higher ploidy is formed solely by duplicating a lower ploidy genome. An example is when a tetraploid is formed by suppression of the first mitotic division in an embryo. This is because the allelic ratios for the higher ploidy are identical to those expected in the lower ploidy. Polyploidy of this kind is relatively rare, and so the method described here is expected to apply in most circumstances.

We suggested a critical value of 10 for the white sturgeon panel based on visual evaluation of the LLR distributions for the known and presumed ploidy white sturgeon. With larger sample sizes, a more quantitative choice of critical values is possible by using those samples to estimate false positive and false negative rates for a given critical value and comparison.

When differentiating between ploidies that are multiples of each other (e.g. diploid and tetraploid), the set of all possible models of the higher ploidy contains all possible models of the lower ploidy. nQuire addresses this for the case of diploids and tetraploids by fixing all three component weights of the tetraploid mixture model at 1/3, effectively assuming that the heterozygous genotype states occur in fixed proportions (Weiß et al., 2018). While they demonstrate the efficacy of this approach, it is unclear whether ploidy inferences would still be accurate when the true genotype proportions have large deviations from those assumed. Gompert and Mock (2017) did not restrict proportions of genotype states. They relied on tetraploids being sufficiently separated from diploids by posterior allele dosages for a clustering algorithm to separate the two categories. The current method also does not restrict proportions of genotype states. As such, when comparing ploidies that are multiples of each other the larger ploidy will always have a likelihood higher than or equal to that of the smaller ploidy (apart from deviations due to the threshold at which convergence is assumed). The larger ploidy can have a higher likelihood due to over-fitting. Ploidy can still be inferred, however, as the distribution of LLR should be approximately bimodal: samples with a true smaller ploidy will have smaller LLR (distributed close to zero), and those with the larger ploidy will have larger LLR (distributed further away from zero). Additionally, the current method estimates the proportion of loci in each genotype state (the component weights) and these can be compared with expectations based on the species’ biology. For example, when comparing diploidy and tetraploidy for a sample and fitting the tetraploid model, if the proportions of genotypes in states 1 and 3 (genotypes of ABBB and AAAB) are estimated to be close to zero, then it may be reasonable to categorize this sample as diploid. This logic is similar to that of restricting the proportions of genotype states.

When integrated into pipelines utilizing amplicon sequencing data, e.g. GT-seq (Campbell et al., 2015), the routine presented herein provides a straightforward and effective method by which samples can be simultaneously genotyped and ploidy inferred from archived as well as fresh tissue samples of diverse types. We provide a convenient R package by which this can be accomplished (tripsAndDipR v 0.2.0 available at www.github.com/delomast/tripsAndDipR). While we encourage users to explore the sequencing depths, heterozygosity, and LLR critical values that provide reliable and robust estimates of ploidy in each particular organism, we expect that this package will empower studies of genetic variation and inheritance in organisms that vary in ploidy naturally or as a result of artificial propagation practices.

## Acknowledgments

We would like to thank staff at Idaho Department of Fish and Game’s Nampa Fish Hatchery, Rapid River Fish Hatchery, and Fish Health Laboratory for providing tissue samples of Chinook salmon and the staff of the Idaho Department of Fish and Game’s Eagle Fish Genetics Laboratory for assistance in genotyping the Chinook salmon samples. We would like to thank Ken Lepla and Idaho Power Company for assistance in acquiring samples. For white sturgeon, Lori Maxwell of CRITFC assisted with collection of laboratory data. Joel Van Eenennaam collected fin clips from known octoploid and dodecaploid white sturgeon. Funding was provided by Bonneville Power Administration project 2008-907-00.

## Author Contributions

TAD derived the statistical model, wrote the R package, assessed the model, and drafted the manuscript. SN and SCW sequenced the white sturgeon samples. AS provided the sturgeon samples and collected the presumed decaploid sturgeon samples. All authors participated in editing and revising the manuscript.

## References

Aegerter, S., & Jalabert, B. (2004). Effects of post-ovulatory oocyte ageing and temperature on egg quality and on the occurrence of triploid fry in rainbow trout, *Oncorhynchus mykiss*. Aquaculture, 231(1-4), 59–71. doi: 10.1016/j.aquaculture.2003.08.019

Augusto Corrêa dos Santos, R., Goldman, G. H., & Riaño-Pachón, D. M. (2017). ploidyNGS: visually exploring ploidy with next generation sequencing data. Bioinformatics, 33(16), 2575–2576. doi: 10.1093/bioinformatics/btx204

Benfey, T. J. (1999). The physiology and behavior of triploid fishes. Reviews in Fisheries Science, 7(1), 39–67. doi: 10.1080/10641269991319162

Byrd, R. H., Lu, P., Nocedal, J., & Zhu, C. (1995). A limited memory algorithm for bound constrained optimization. SIAM Journal on Scientific Computing, 16(5), 1190–1208. doi: 10.1137/0916069

Campbell, N. R., Harmon, S. A., & Narum, S. R. (2015). Genotyping-in-Thousands by sequencing (GT-seq): A cost effective SNP genotyping method based on custom amplicon sequencing. Molecular Ecology Resources, 15(4), 855–867. doi: 10.1111/1755-0998.12357

Cassinelli, J. D., Meyer, K. A., Koenig, M. K., Vu, N. V, & Campbell, M. R. (2018). Performance of diploid and triploid westslope cutthroat trout fry stocked into Idaho alpine lakes. North American Journal of Fishery Management, 39(1), 112–123. doi: 10.1002/nafm.10254

Cherfas, N., Gomelsky, B., Ben-Dom, N., & Hulata, G. (1995). Evidence for the heritable nature of spontaneous diploidization in common carp, *Cyprinus carpio* L., eggs. Aquaculture Research, 26(4), 289–292. doi: 10.1111/j.1365-2109.1995.tb00914.x

Delomas, T. A. (2019). Differentiating diploid and triploid individuals using single nucleotide polymorphisms genotyped by amplicon sequencing. Molecular Ecology Resources, 19(6), 1545–1551. doi: 10.1111/1755-0998.13073

Delomas, T. A., & Dabrowski, K. (2016). Zebrafish embryonic development is induced by carp sperm. Biology Letters, 12(11), 20160628.

Delomas, T. A., & Dabrowski, K. (2018). Why are triploid zebrafish all male? Molecular Reproduction and Development, 85(7), 612–621. doi: 10.1002/mrd.22998

Drauch Schreier, A., Gille, D., Mahardja, B., & May, B. (2011). Neutral markers confirm the octoploid origin and reveal spontaneous autopolyploidy in white sturgeon, *Acipenser transmontanus*. Journal of Applied Ichthyology, 27(SUPPL. 2), 24–33. doi: 10.1111/j.1439-0426.2011.01873.x

Feindel, N. J., Benfey, T. J., & Trippel, E. A. (2010). Competitive spawning success and fertility of triploid male Atlantic cod *Gadus morhua*. Aquaculture Environment Interactions, 1, 47–55. doi: 10.2307/24864017

Fiske, J. A., Van Eenennaam, J. P., Todgham, A. E., Young, S. P., Holem-Bell, C. E., Goodbla, A. M., & Schreier, A. D. (2019). A comparison of methods for determining ploidy in white sturgeon *(Acipenser transmontanus)*. Aquaculture, 507, 435–442. doi: 10.1016/j.aquaculture.2019.03.009

Flajšhans, M., Kvasnička, P., & Ráb, P. (1993). Genetic studies in tench (*Tinca tinca* L.): high incidence of spontaneous triploidy. Aquaculture, 110(3–4), 243–248. doi: 10.1016/0044-8486(93)90372-6

Gerard, D., Ferrão, L. F. V., Garcia, A. A. F., & Stephens, M. (2018). Genotyping polyploids from messy sequencing data. Genetics, 210(3), 789–807. doi: 10.1534/genetics.118.301468

Glover, K. A., Madhun, A. S., Dahle, G., Sørvik, A. G. E., Wennevik, V., Skaala, Ø.,… Fjelldal, P. G. (2015). The frequency of spontaneous triploidy in farmed Atlantic salmon produced in Norway during the period 2007–2014. BMC Genetics, 16(1), 37. doi: 10.1186/s12863-015-0193-0

Gold, J. R., & Avise, J. C. (1976). Spontaneous triploidy in the California roach *Hesperoleucus symmetricus* (Pisces: Cyprinidae). Cytogenetic and Genome Research, 17(3), 144–149. doi: 10.1159/000130706

Gompert, Z., & Mock, K. E. (2017). Detection of individual ploidy levels with genotyping-by-sequencing (GBS) analysis. Molecular Ecology Resources, 17(6), 1156–1167. doi: 10.1111/1755-0998.12657

Husband, B. C., & Sabara, H. A. (2004, March 1). Reproductive isolation between autotetraploids and their diploid progenitors in fireweed, *Chamerion angustifolium* (Onagraceae). New Phytologist, Vol. 161, pp. 703–713. doi: 10.1046/j.1469-8137.2004.00998.x

Husband, B. C., Schemske, D. W., Burton, T. L., & Goodwillie, C. (2002). Pollen competition as a unilateral reproductive barrier between sympatric diploid and tetraploid *Chamerion angustifolium*. Proceedings of the Royal Society of London. Series B: Biological Sciences, 269(1509), 2565–2571. doi: 10.1098/rspb.2002.2196

Hyndman, C. A., Kieffer, J. D., & Benfey, T. J. (2003). Physiology and survival of triploid brook trout following exhaustive exercise in warm water. Aquaculture, 221(1–4), 629–643. doi: 10.1016/S0044-8486(03)00119-4

Lamatsch, D. K., & Stöck, M. (2009). Sperm-dependent parthenogenesis and hybridogenesis in teleost fishes. In I. Schön, K. Martens, & P. Dijk (Eds.), Lost Sex (pp. 399–432). doi: 10.1007/978-90-481-2770-2_19

Leal, M. J., Clark, B. E., Van Eenennaam, J. P., Schreier, A. D., & Todgham, A. E. (2018). The effects of warm temperature acclimation on constitutive stress, immunity, and metabolism in white sturgeon (*Acipenser transmontanus*) of different ploidies. Comparative Biochemistry and Physiology-Part A□: Molecular and Integrative Physiology, 224, 23–34. doi: 10.1016/j.cbpa.2018.05.021

Liu, S., Liu, Y., Zhou, G., Zhang, X., Luo, C., Feng, H.,… Yang, H. (2001). The formation of tetraploid stocks of red crucian carp × common carp hybrids as an effect of interspecific hybridization. Aquaculture, 192(2), 171–186. doi: 10.1016/S0044-8486(00)00451-8

Machado, S. N., Neto, M. F., Bakkali, M., Vicari, M. R., Artoni, R. F., Oliveira, C. de, & Foresti, F. (2012). Natural triploidy and B chromosomes in *Astyanax scabripinnis* (Characiformes, Characidae): a new occurrence. Caryologia, 65(1), 40–46. doi: 10.1080/00087114.2012.678086

Mock, K. E., Callahan, C. M., Islam-Faridi, M. N., Shaw, J. D., Rai, H. S., Sanderson, S. C.,… Wolf, P. G. (2012). Widespread triploidy in western North American aspen *(Populus tremuloides)*. PLoS ONE, 7(10), e48406. doi: 10.1371/journal.pone.0048406

Nell, J. A. (2002). Farming triploid oysters. Aquaculture, 210(1-4), 69–88. doi: 10.1016/S0044-8486(01)00861-4

Ptacek, M. B., Gerhardt, H. C., & Sage, R. D. (1994). Speciation by polyploidy in treefrogs: Multiple origins of the tetraploid, *Hyla versicolor*. Evolution, 48(3), 898–908. doi: 10.1111/j.1558-5646.1994.tb01370.x

Thorgaard, G. H., Rabinovitch, P. S., Shen, M. W., Gall, G. A. E., Propp, J., & Utter, F. M. (1982). Triploid rainbow trout identified by flow cytometry. Aquaculture, 29(3–4), 305–309. doi: 10.1016/0044-8486(82)90144-2

Utsunomia, R., Pansonato Alves, J. C., Paiva, L. R. S., Costa Silva, G. J., Oliveira, C., Bertollo, L. A. C., & Foresti, F. (2014). Genetic differentiation among distinct karyomorphs of the wolf fish *Hoplias malabaricus* species complex (Characiformes, Erythrinidae) and report of unusual hybridization with natural triploidy. Journal of Fish Biology, 85(5), 1682–1692. doi: 10.1111/jfb.12526

Van Eenennaam, J. P., Fiske, A. J., Leal, M. J., Cooley-Rieders, C., Todgham, A. E., Conte, F. S., & Schreier, A. D. (2020). Mechanical shock during egg de-adhesion and post-ovulatory ageing contribute to spontaneous autopolyploidy in white sturgeon culture *(Acipenser transmontanus)*. Aquaculture, 515, 734530. doi: 10.1016/j.aquaculture.2019.734530

Wattendorf, R. J. (1986). Rapid identification of triploid grass carp with a Coulter counter and channelyzer. The Progressive Fish□Culturist, 48(2), 125–132. doi: 10.1577/1548-8640(1986)48<125:RIOTGC>2.0.CO;2

Weiß, C. L., Pais, M., Cano, L. M., Kamoun, S., & Burbano, H. A. (2018). nQuire: a statistical framework for ploidy estimation using next generation sequencing. BMC Bioinformatics, 19(1), 122. doi: 10.1186/s12859-018-2128-z

Willis, S. C., Delomas, T. A., Parker, B., Miller, D., Anders, P., & Narum, S. (2020). Single nucleotide polymorphism genotypes and ploidy estimates for ploidy variable species generated with massively parallel amplicon sequencing. Preprint/Submitted, 0.

Wood, T. E., Takebayashi, N., Barker, M. S., Mayrose, I., Greenspoon, P. B., & Rieseberg, L. H. (2009). The frequency of polyploid speciation in vascular plants. Proceedings of the National Academy of Sciences of the United States of America, 106(33), 13875–13879. doi: 10.1073/pnas.0811575106

Yamashita, M., Jiang, J., Onozato, H., Nakanishi, T., & Nagahama, Y. (1993). A tripolar spindle formed at meiosis I assures the retention of the original ploidy in the gynogenetic triploid crucian carp, ginbuna *Carassius auratus langsdorfii*. Develop. Growth & Differ, 35(6), 631–636. doi: 10.1111/j.1440-169X.1993.00631.x

Zhang, Q., & Arai, K. (1999). Distribution and reproductive capacity of natural triploid individuals and occurrence of unreduced eggs as a cause of polyploidization in the loach, *Misgurnus anguillicaudatus*. Ichthyological Research, 46(2), 153–161. doi: 10.1007/BF02675433

